# Influence of Mpv17 on hair-cell mitochondrial homeostasis, synapse integrity, and vulnerability to damage in the zebrafish lateral line

**DOI:** 10.1101/2021.04.12.439521

**Authors:** Melanie Holmgren, Lavinia Sheets

## Abstract

Noise exposure is particularly stressful to hair-cell mitochondria, which must produce enough energy to meet high metabolic demands as well as regulate local intracellular Ca^2+^ concentrations. Mitochondrial Inner Membrane Protein 17 (Mpv17) functions as a non-selective channel and plays a role in maintaining mitochondrial homeostasis. In zebrafish, hair cells in *mpv17^a9/a9^* mutants displayed elevated levels of reactive oxygen species (ROS), elevated mitochondrial calcium, hyperpolarized transmembrane potential, and greater vulnerability to neomycin, indicating impaired mitochondrial function. Using a strong water current to overstimulate hair cells in the zebrafish lateral line, we observed *mpv17^a9/a9^* mutant hair cells were more vulnerable to morphological disruption and hair-cell loss than wild type siblings simultaneously exposed to the same stimulus. To determine the role of mitochondrial homeostasis on hair-cell synapse integrity, we surveyed synapse number in *mpv17^a9/a9^* mutants and wild type siblings as well as the sizes of presynaptic dense bodies (ribbons) and postsynaptic densities immediately following stimulus exposure. We observed mechanically injured *mpv17^a9/a9^* neuromasts, while they lost a greater number of hair cells, lost a similar number of synapses per hair cell relative to wild type. Additionally, we quantified the size of hair cell pre- and postsynaptic structures and observed significantly enlarged wild type postsynaptic densities, yet relatively little change in the size of *mpv17^a9/a9^* postsynaptic densities following stimulation. These results suggest impaired hair-cell mitochondrial activity influences synaptic morphology and hair-cell survival but does not exacerbate synapse loss following mechanical injury.

## Introduction

Hair cells, the sensory receptors of the inner ear, rely on mitochondria to generate energy to meet the high metabolic demands of mechanotransduction and synaptic transmission (Puschner & Schacht, 1997). Hair-cell mitochondria also produce reactive oxygen species (ROS) and contribute to the homeostatic control of intracellular Ca^2+^ (Collins et al., 2012; Matlib et al., 1998; Wong et al., 2019; Zenisek & Matthews, 2000). Disruption of mitochondrial dynamics can affect hair-cell vulnerability to damage from ototoxic drugs, as well as interfere with maintenance of hair-cell synapses (Esterberg, Hailey, Rubel, & Raible, 2014; Esterberg et al., 2016; Wong et al., 2019).

Zebrafish have emerged as a powerful tool to study the roles of mitochondria in hair-cell damage (Holmgren & Sheets, 2020). In addition to their ears, zebrafish have hair cells in their lateral line organs. Composed of clusters of innervated hair cells and supporting cells called neuromasts, zebrafish use their lateral line organs to detect local water currents and mediate behaviors such as escape responses and rheotaxis (Dijkgraaf, 1963; Olive et al., 2016; Olszewski, Haehnel, Taguchi, & Liao, 2012; Stewart, Cardenas, & McHenry, 2013; Suli, Watson, Rubel, & Raible, 2012). Unlike cochlear hair cells, lateral-line hair cells are superficially located on the surface of the body and are therefore pharmacologically and optically accessible in an intact fish. Zebrafish became established as a model system for studying hair-cell development and function due to identification of numerous conserved genes involved in hearing and balance (Nicolson, 2017).

Hearing loss is common in patients with mitochondrial disorders (Hsu et al., 2005). One gene that has been associated with mitochondrial disease in mammals and is conserved in zebrafish is *mpv17* (Krauss et al., 2013; Muller et al., 1997). *mpv17* encodes Mitochondrial Inner Membrane Protein 17, which is a non-selective cation channel that modulates the mitochondrial potential and contributes to mitochondrial homeostasis (Antonenkov et al., 2015; Jacinto et al., 2021). Mice lacking Mpv17 show severe defects in the kidney and inner ear (Meyer zum Gottesberge, Felix, Reuter, & Weiher, 2001; Meyer zum Gottesberge, Massing, & Hansen, 2012; Muller et al., 1997). In contrast, zebrafish lacking Mpv17 appear healthy and have normal life spans. Two zebrafish lines containing a spontaneous mutation in *mpv17* (*roy orbison* (*mpv17^a9/a9^*) and *transparent* (*mpv17^b18/b18^*)) both contain the same 19 bp deletion leading to aberrant splicing and a premature stop codon (D’Agati et al., 2017; Krauss et al., 2013). Notably, the *mpv17^a9/a9^* mutation is carried in the Casper strain of zebrafish, which are commonly used for imaging studies because they lack iridophores and thus have transparent skin (Martorano et al., 2019; White et al., 2008). Mpv17 has been shown in zebrafish to localize to mitochondria in multiple cell types, including lateral-line hair cells (Krauss et al., 2013). Although Casper fish are commonly used in research, how loss of Mpv17 affects mitochondrial function in hair cells of the zebrafish lateral line has not yet been characterized. Additionally, as mitochondrial dysfunction is known to contribute to the pathologies underlying noise-induced hearing loss (Bottger & Schacht, 2013), we further wanted to examine the role of mitochondria homeostasis in mechanically induced hair-cell damage.

In this study, we investigated how loss of Mpv17 affects mitochondrial function in zebrafish lateral line hair cells as well as vulnerability to mechanical injury. In *mpv17^a9/a9^* hair cells, we observed elevated ROS and mitochondrial Ca^2+^, reduced FM1-43 uptake, and increased sensitivity to neomycin-induced hair-cell death. We have previously reported a protocol using a strong water current stimulus to induce mechanical damage to zebrafish lateral-line organs (Holmgren et al., 2021). When exposed to the same stimulus as wild type siblings, mechanically overstimulated *mpv17^a9/a9^* neuromasts were more vulnerable to morphological disruption and hair-cell loss but showed similar degrees of de-innervation and synapse loss. Our results suggest that genetic disruption of mitochondrial homeostasis influences vulnerability to ototoxic or mechanically induced hair-cell death but does not exacerbate mechanically induced hair-cell synapse loss.

## Results

### Mitochondrial homeostasis is disrupted in *mpv17^a9/a9^* hair cells

It has been previously reported that zebrafish lacking Mpv17 have impaired mitochondrial function and that Mpv17 protein localizes to hair-cell mitochondria (Krauss et al., 2013; Martorano et al., 2019). We thus characterized how loss of Mpv17 affected mitochondrial homeostasis in hair cells of the lateral line. To quantify ROS levels in lateral-line hair cells, we exposed *mpv17^a9/a9^* larvae and their wild type siblings to the probe CellROX Green (Fig. 1 A,B). We observed increased fluorescence in *mpv17^a9/a9^* neuromasts relative to wild type, indicating elevated ROS in *mpv17^a9/a9^* hair cells (Fig. 1 C; Mann-Whitney test **** P<0.0001). We then quantified baseline mitochondrial Ca^2+^ levels using the genetically encoded indicator mitoGCaMP3 (Fig. 1 D,E) (Esterberg et al., 2014). Relative to wild type siblings, *mpv17^a9/a9^* neuromasts had increased mitoGCaMP3 fluorescence, indicative of elevated mitochondrial Ca^2+^(Fig. 1 F; Mann-Whitney test ****P<0.0001).

**Figure 1.**
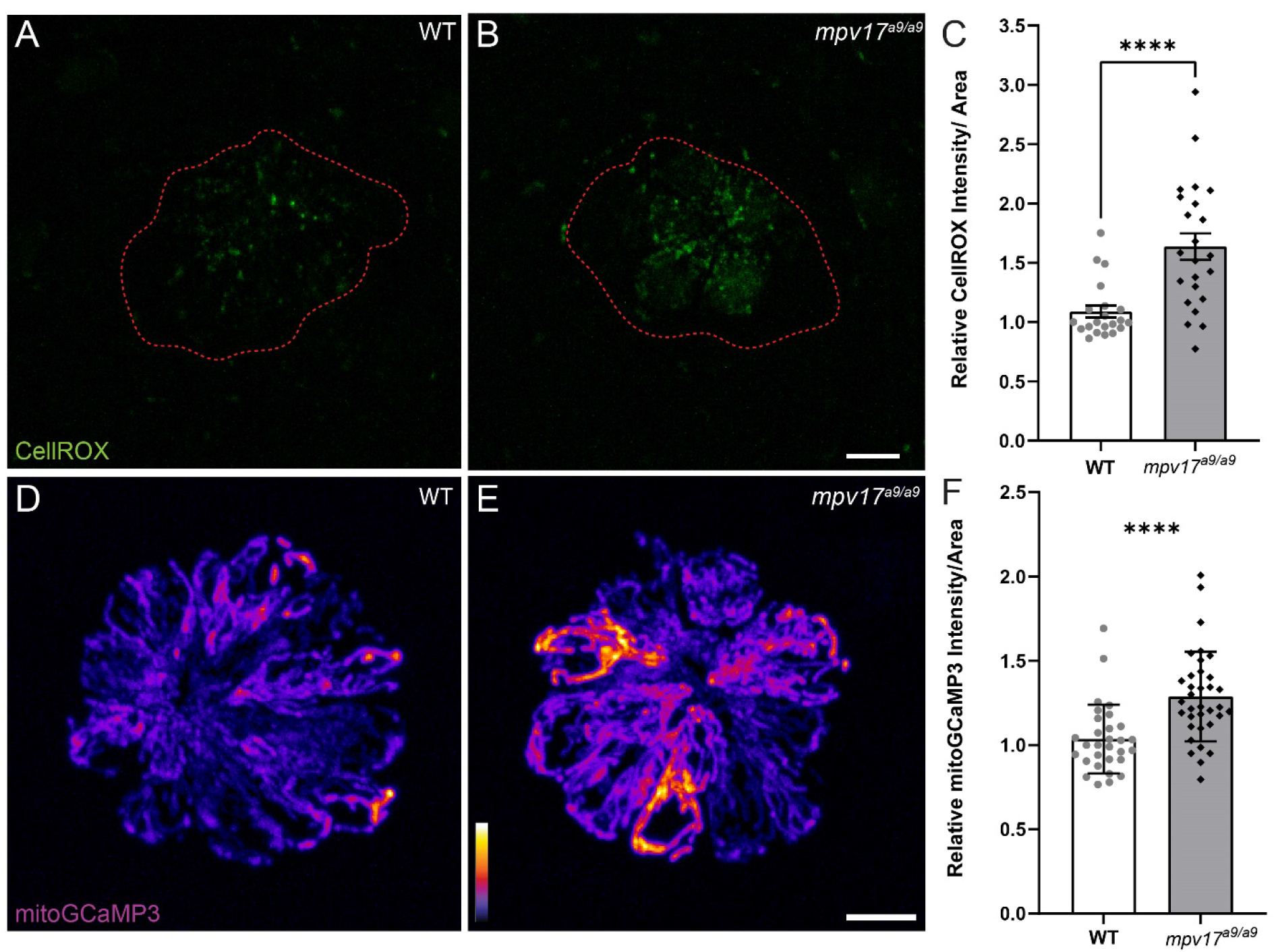
Mitochondrial homeostasis in wild type and *mpv17^a9/a9^* neuromasts. **(A-B)**Maximum-intensity projections of confocal images showing CellROX Green staining in wild type siblings (WT) (A) and *mpv17^a9/a9^* (B) neuromasts. Neuromast boundaries were delineated based on DIC images (not shown). **(C)** Mean CellROX intensity is elevated in *mpv17^a9/a9^* neuromasts (****P<0.0001). **(D-E)** Maximum-intensity projections of confocal images showing mitoGCaMP3 fluorescence in WT (D) and *mpv17^a9/a9^* (E) neuromasts. **(F)** Mean mitoGCaMP3 intensity is increased in *mpv17^a9/a9^* neuromasts (****P<0.0001). Scale bars: 5 μm.

It has been shown in murine fibroblasts that loss of Mpv17 results in hyperpolarized mitochondria (Antonenkov et al., 2015). To measure mitochondrial membrane potential (ΔΨ_m_) of lateral-line hair cells, we exposed *mpv17^a9/a9^* and sibling larvae to the dyes MitoTracker Red CMXRos and MitoTracker Deep Red (Fig. 2 B,C,F,G). Both of these dyes are well-retained after fixation and their accumulation in mitochondria is dependent on ΔΨ_m_ (Mot, Liddell, White, & Crouch, 2016; Pendergrass, Wolf, & Poot, 2004). We verified that these probes could be used to detect hyperpolarized ΔΨ_m_ by treating wild type larvae with cyclosporin A (Supplemental Fig. 1 A,B). We also verified that MitoTracker entry into hair cells is not dependent on hair-cell mechanotransduction by briefly treating larvae with BAPTA to disrupt tip links prior to MitoTracker exposure (Supplemental Fig. 1 A,B). With both probes, we observed increased fluorescence in *mpv17^a9/a9^* neuromasts relative to wild type, indicating *mpv17^a9/a9^* hair cells have hyperpolarized mitochondria (Fig. 2 D, H; *P=0.0469 MitoTracker Red CMXRos; ****P<0.0001 MitoTracker Deep Red).

**Figure 2.**
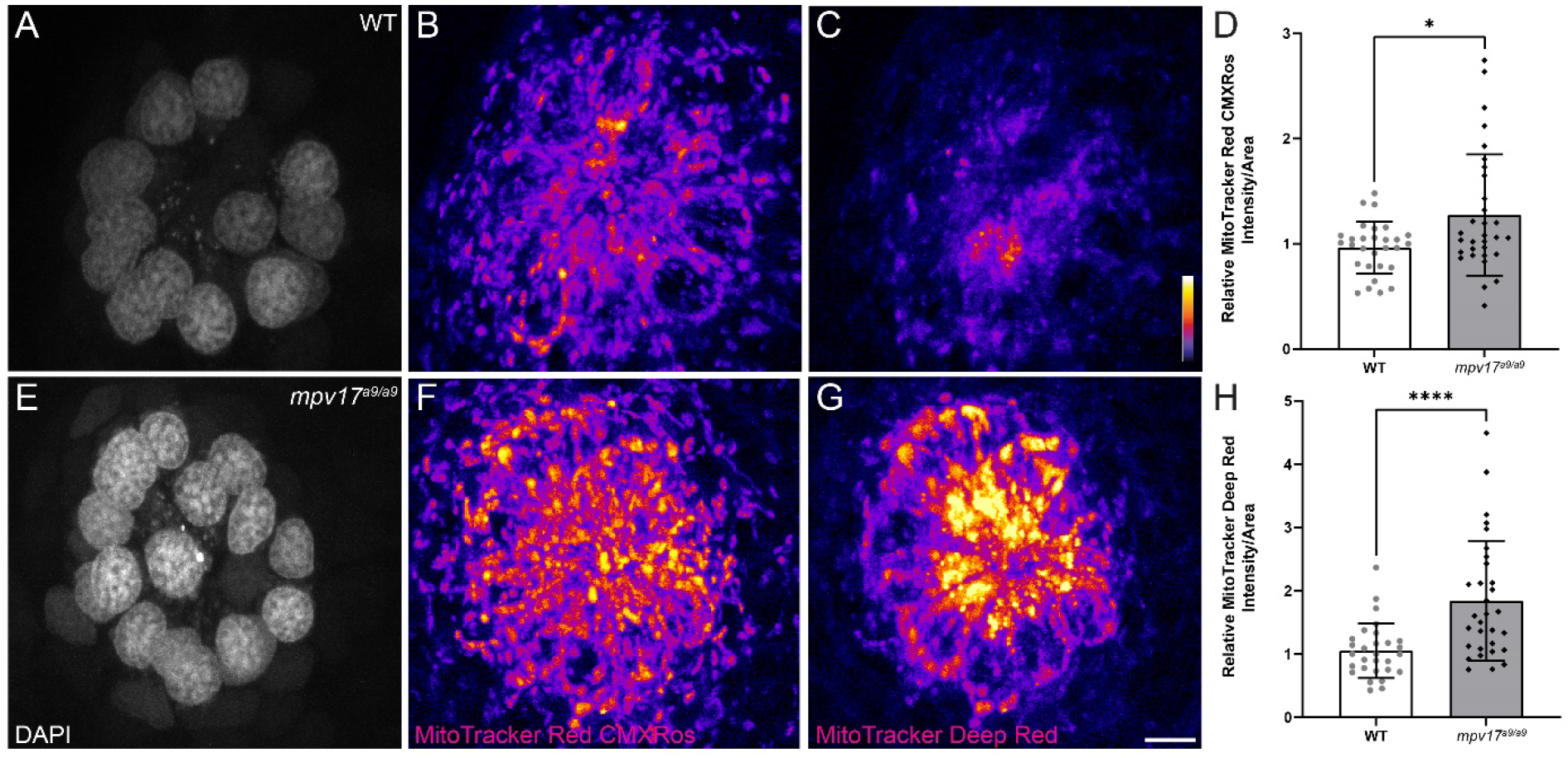
*mpv17^a9/a9^* hair cells have hyperpolarized mitochondria. **(A-C, E-G)** Maximum-intensity projections of confocal images displaying wild type (A-C) and *mpv17^a9/a9^* (EG) neuromasts with DAPI-labeled hair cells (A,E) and staining with MitoTracker Red CMXRos (B,F) and MitoTracker Deep Red (C,G). **(D, H)** Mean intensities for both MitoTracker Red CMXRos (D) and MitoTracker Deep Red (H) are elevated in *mpv17^a9/a9^* neuromasts (*P=0.0469; ****P<0.0001). Scale bar: 5 μm.

To determine the effect of Mpv17 deficiency on hair-cell function, we next exposed *mpv17^a9/a9^* and wild type larvae to the vital dye FM1-43FX (Fig. 3 A,B). Labeling of hair cells following brief exposure to FM1-43 occurs via active mechanotransduction, and reduced labeling indicates reduced driving force for cations through mechanotransduction channels (Toro et al., 2015). We observed significantly reduced uptake of this dye in *mpv17^a9/a9^* neuromasts relative to wild type (Fig. 3 C; Unpaired t test ***P=0.0002), suggesting impaired mechanotransduction in *mpv17^a9/a9^* hair cells. Collectively, these results support that mitochondrial homeostasis is disrupted in *mpv17^a9/a9^* hair cells, and that hair-cell transduction is also reduced.

**Figure 3.**
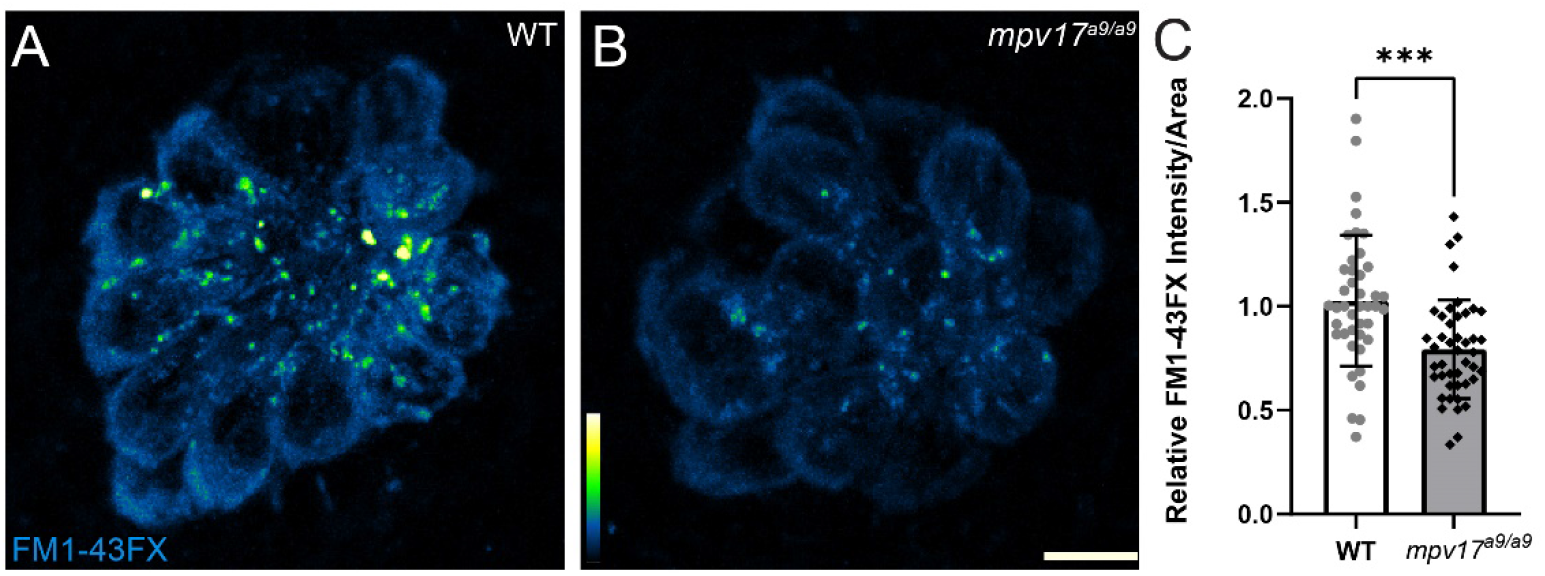
FM1-43 uptake is reduced in *mpv17^a9/a9^* hair cells. **(A-B)** Maximum-intensity projections of confocal images showing wild type (A) and *mpv17^a9/a9^* (B) neuromasts exposed to FM1-43FX. **(C)** Mean intensity of FM1-43FX is reduced in *mpv17^a9/a9^* neuromasts (***P=0.0002). Scale bar: 5 μm.

### *mpv17^a9/a9^* hair cells are more susceptible to neomycin-induced death

A recent study demonstrated that zebrafish mutants with elevated ROS are more vulnerable to neomycin-induced hair-cell loss (Alassaf, Daykin, Mathiaparanam, & Wolman, 2019). It has also been shown that elevated mitochondrial Ca^2+^ or ΔΨ_m_ increases sensitivity to neomycin (Esterberg et al., 2014). On the other hand, it is known that neomycin enters hair cells through mechanotransduction channels, and blocking mechanotransduction can reduce neomycin sensitivity (Alharazneh et al., 2011; Hailey, Esterberg, Linbo, Rubel, & Raible, 2017; Owens et al., 2009). We therefore exposed *mpv17^a9/a9^* and wild type sibling larvae to a low dose of neomycin to determine whether *mpv17^a9/a9^* larvae are more or less sensitive to neomycin-induced hair-cell death. Both groups lost a significant number of hair cells following exposure to 10 μM neomycin (Fig. 4; Repeated measure two-way ANOVA ****P<0.0001 wild type; ****P<0.0001 *mpv17^−/−^*), however this loss was significantly more severe in *mpv17^a9/a9^* larvae (Repeated measure two-way ANOVA ***P=0.0007). Thus, loss of Mpv17 increases sensitivity to neomycin-induced hair-cell death.

**Figure 4.**
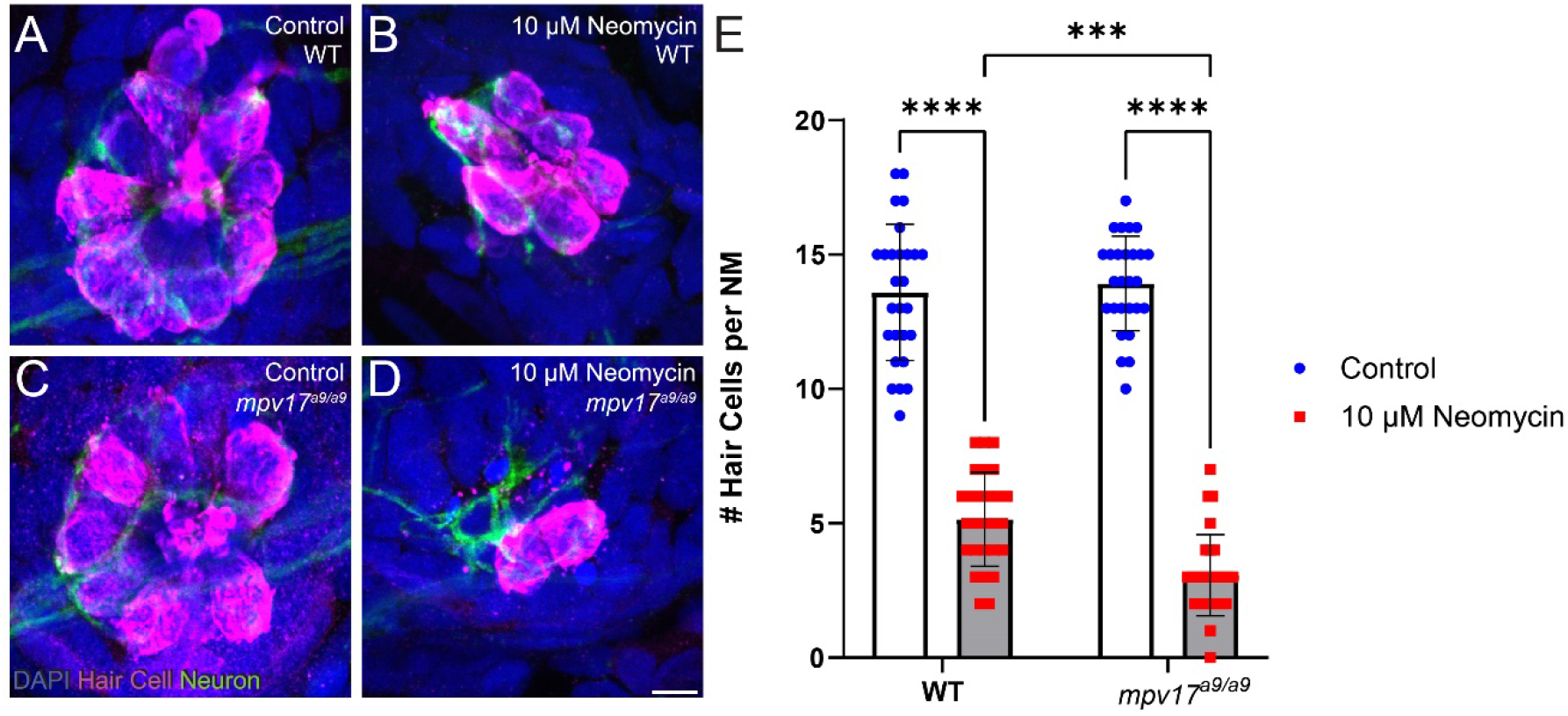
*mpv17^a9/a9^* larvae are more sensitive to neomycin-induced hair-cell loss. **(A-D)** Maximum-intensity projections of confocal images of wild type (A,B) and *mpv17^a9/a9^* (C,D) control neuromasts (A,C) or exposed to 10 μM neomycin (B,D). Hair cells were visualized with immunolabel of Parvalbumin (magenta), afferent neurites are expressing GFP (green), and all cell nuclei are labeled with DAPI (blue). **(E)** Number of hair cells per neuromasts was significantly reduced in neuromasts exposed to 10 μM neomycin, but *mpv17^a9/a9^* neuromasts were significantly more sensitive than wild type (****P<0.0001, ***P=0.0007). Scale bar: 5 μm.

### *mpv17^a9/a9^* hair cells are more vulnerable to mechanically-induced morphological disruption and hair-cell loss

We have previously reported a protocol to mechanically overstimulate zebrafish lateral line organs using strong water current (Holmgren et al., 2021). This stimulation resulted in phenotypes including mechanical disruption of neuromast morphology, loss of hair cells, neurite retraction, and loss of hair-cell synapses. To determine the effects of impaired mitochondrial homeostasis on mechanically induced lateral-line damage, we exposed 7-day-old *mpv17^a9/a9^* larvae and wild type siblings to strong water current, then fixed them for immunohistochemical labeling of hair cells, neurites, and synaptic components (Fig. 5 A-C). As indicated in our previous study, image analysis was performed on blinded samples. As in our previous study, in fish exposed to strong water current we observed two distinct morphological profiles of the neuromasts: “normal” in which the neuromasts appeared identical to controls with the hair cells arranged in a typical rosette structure (Fig. 5 B), or “disrupted,” in which the neuromasts were displaced on their sides with elongated and misshapen hair cells and the apical ends of the hair cells oriented posteriorly (Fig 5 C). In stimulus exposed wild type fish, we observed disrupted morphology in 41% of the neuromasts surveyed. We observed this morphological change more frequently in the more posterior L5 neuromasts compared to the more anterior L3 neuromasts (Fig. 5 E; 56% L5; 39% L4; 27% L3). We observed in *mpv17^a9/a9^* larvae a similar trend of increased morphological disruption in the more posterior neuromasts, however the frequency of disrupted neuromasts was higher in mechanically injured *mpv17^a9/a9^* larvae compared to wild type siblings concurrently exposed to the same stimulus (75% L5; 69% L4; 52% L3). Thus, *mpv17^a9/a9^* neuromasts appear to be more vulnerable to morphological disruption resulting from mechanical injury.

**Figure 5.**
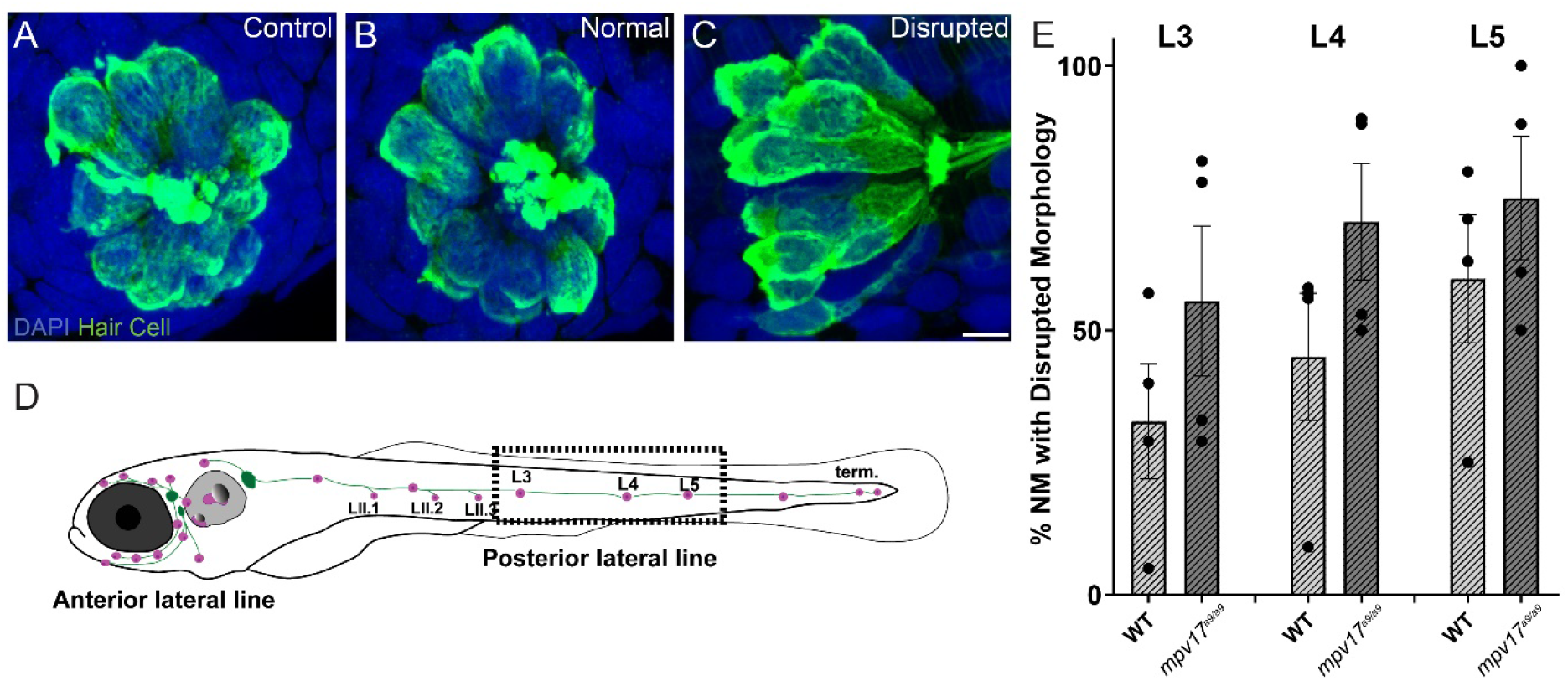
Mechanical injury results in morphological disruption of neuromasts more frequently in *mpv17^a9/a9^* larvae than in wild type. **(A-C)** Maximum-intensity projections of confocal images showing hair cells labeled with Parvalbumin and all nuclei labeled with DAPI in control neuromasts (A) and strong water current exposed neuromasts with normal (B) or disrupted (C) morphology. **(D)** Schematic of larval zebrafish indicating the placement of neuromasts (pink dots) and afferent nerves (green lines). Neuromasts L3-5 (dashed rectangle) were examined in this study. **(E)** Quantification of disrupted neuromasts. Each point indicates the percentage of neuromasts with disrupted morphology in a single experimental trial. The frequency of disrupted neuromasts was higher in the more posterior L5 neuromasts, *mpv17^a9/a9^* neuromasts showed disrupted morphology more frequently than wild type. Scale bar: 5 μm.

We then compared hair-cell numbers between mechanically injured *mpv17^a9/a9^* and wild type neuromasts. Wild type sibling fish exhibited a slight decrease in the number of hair cells per neuromast (Fig. 6 G; Repeated measure two-way ANOVA P=0.0791). Similarly, *mpv17^a9/a9^* larvae exposed to the same stimulus showed a significant decrease in hair-cell number (Repeated measure two-way ANOVA *P=0.0117). In both groups, this reduction in hair-cell number was specific to disrupted neuromasts; hair-cell numbers in exposed neuromasts with “normal” hair-cell morphology were comparable to control (Fig. 6 H; Repeated measure two-way ANOVA P=0.9624 wild type normal; P=0.0180 wild type disrupted; P>0.9999 *mpv17^a9/a9^* normal; ****P<0.0001 *mpv17^a9/a9^* disrupted). We also observed a significant reduction in the percentage of hair cells per neuromast with *neurod:EGFP* labeled afferent contacts in both wild type and *mpv17^a9/a9^* neuromasts (Fig. 6 I; One sample Wilcoxon test ***P=0.0002 wild type; ***P=0.0005 *mpv17^a9/a9^*). Similar to the reduction in hair-cell number, this neurite retraction phenotype was evident only in “disrupted” neuromasts of both groups (Fig. 6 J; One sample Wilcoxon test ***P=0.0002 wild type disrupted; ***P=0.0005 *mpv17^a9/a9^* disrupted). Taken together, these results demonstrate that *mpv17^a9/a9^* neuromasts are more susceptible to mechanically induced morphological disruption and hair-cell loss, but that impaired mitochondrial function does not affect sensitivity to afferent retraction resulting from mechanical injury.

**Figure 6.**
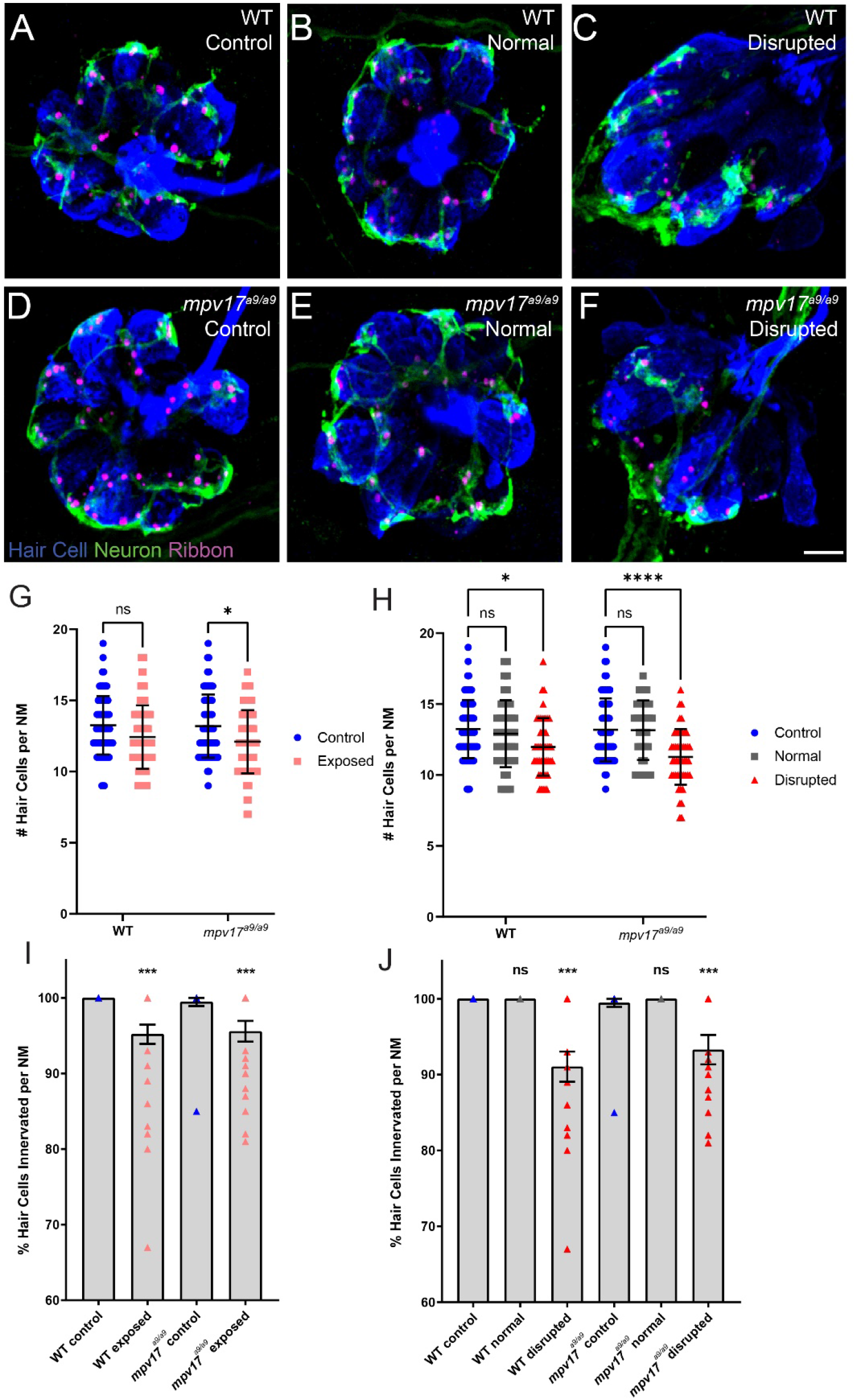
Wild type and *mpv17 ^a9/a9^* neuromasts show loss of hair cells and afferent innervation following mechanical overstimulation. **(A-F)** Maximum-intensity projections of confocal images showing neuromasts with Parvalbumin-labeled hair cells (blue) and Ribeye b-labeled presynaptic ribbons (magenta). Neurod:GFP-labeled afferent neurites are also shown (green). Unexposed control neuromasts are shown in (A; wild type (WT)) and (D; *mpv17^a9/a9^*); exposed neuromasts with normal morphology are shown in (B; WT) and (E; *mpv17^a9/a9^*); and exposed neuromasts with disrupted morphology are shown in (C; WT) and (F; *mpv17^a9/a9^*). **(G-H)** Quantification of hair cells per neuromast shows a significant loss of hair cells in exposed *mpv17 ^a9/a9^*-neuromasts (G; *P=0.0117), which is specific to disrupted neuromasts (H; *P=0.0180; ****P<0.0001). **(I-J)** Percentage of hair cells with GFP-labeled contacts. We observed significant neurite retraction (I; ***P=0.0002 (WT); ***P=0.0005 (*mpv17 ^a9/a9^*)), which is also specific to disrupted neuromasts only (J; ***P=0.0002 (WT); ***P=0.0005 (*mpv17^a9/a9^*)). Scale bar: 5 μm.

### Wild type and *mpv17^a9/a9^* hair cells show comparable synapse loss following mechanical overstimulation

Recent studies indicate hair-cell mitochondrial activity plays a key role in synaptic maintenance (Wang et al., 2018; Wong et al., 2019). We have previously shown that mechanical overstimulation resulted in loss of hair-cell synapses, as well as changes in synapse morphology (Holmgren et al., 2021). To determine the effect of impaired mitochondrial function on synapse number following mechanical overstimulation, we exposed *mpv17^a9/a9^* fish and their wild type siblings, then immunolabeled synaptic components and quantified intact synapses, defined as presynaptic ribbons colocalized with postsynaptic densities (PSD) (Fig. 7 A-F). In mechanically overstimulated wild type neuromasts, we observed a statistically significant decrease in the number of intact synapses per hair cell (Fig. 7 G; Repeated measure two-way ANOVA **P=0.0016 wild type). In agreement with our previous study, this loss of synapses occurred in all exposed neuromasts i.e. both normal and disrupted morphologies (Repeated measure two-way ANOVA **P=0.0090 wild type normal; **P=0.0079 wild type disrupted), suggesting that synapse loss may be a consequence of hair-cell overstimulation rather than physical mechanical injury. In mechanically overstimulated *mpv17^a9/a9^* fish, we observed a similar reduction in synapse number (Repeated measure twoway ANOVA **P=0.0014), and this trend of synapse loss occurred in all exposed neuromasts (Repeated measure two-way ANOVA P=0.0948 *mpv17^a9/a9^* normal; P=0.0505 *mpv17^a9/a9^* disrupted). The similarity in synapse loss between mutants and wild type siblings suggest loss of Mpv17 does not dramatically affect mechanically induced synapse loss. Also notable was that unexposed control wild type and *mpv17^a9/a9^* fish had a comparable number of synapses per hair cell (Fig. 7 G; P=0.9973) indicating that chronic mitochondrial disfunction in *mpv17^a9/a9^*hair cells does not affect synaptic maintenance.

**Figure 7.**
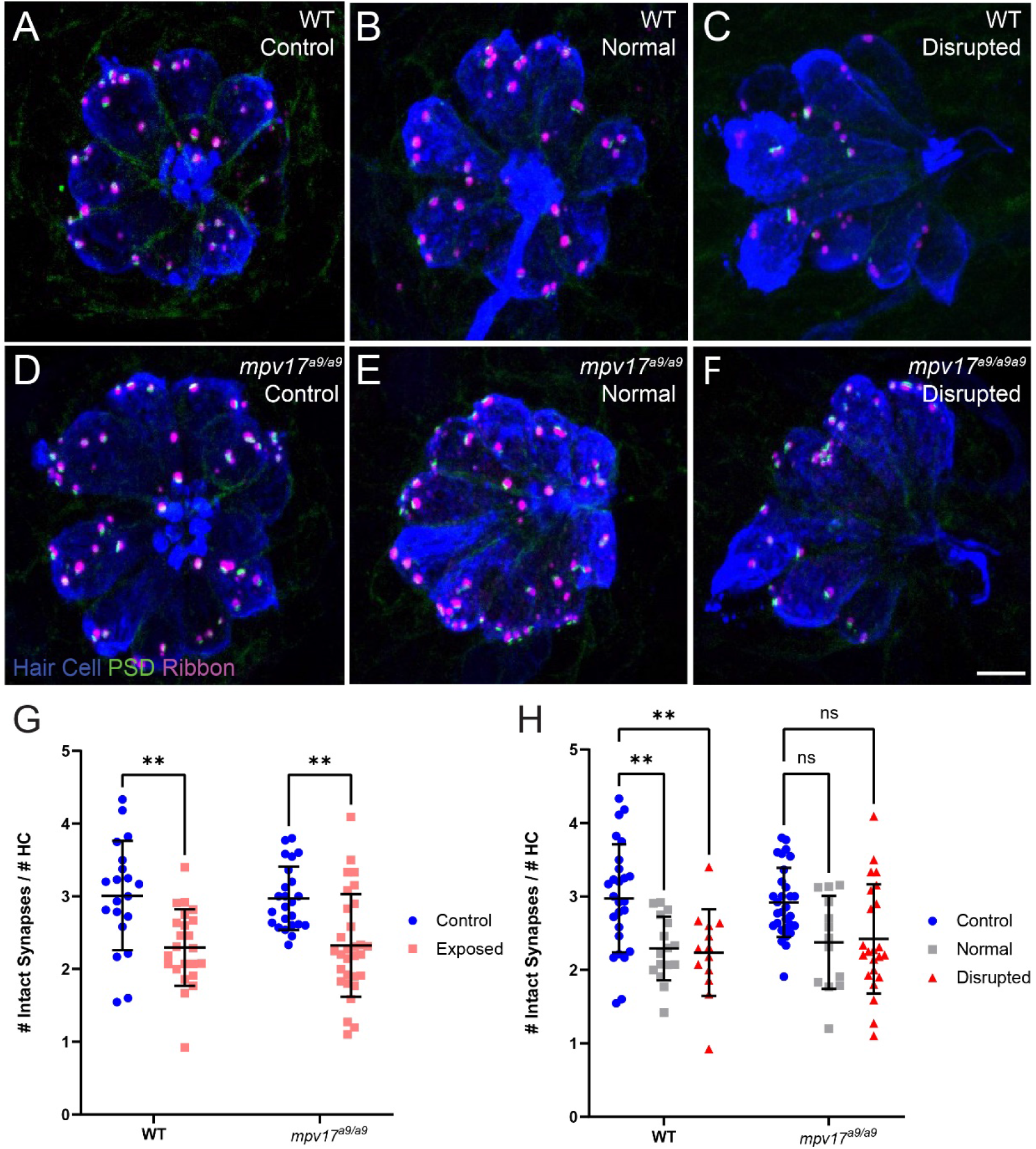
Both wild type and *mpv17^a9/a9^* neuromasts experience mechanically induced hair-cell synapse loss. **(A-C)** Maximum-intensity projections of confocal images showing neuromasts with Parvalbumin-labeled hair cells (blue), Ribeye b-labeled presynaptic ribbons (magenta), and MAGUK-labeled PSD (green). Unexposed control neuromasts are shown in (A; wild type (WT)) and (D; *mpv17^a9/a9^*); exposed neuromasts with normal morphology are shown in (B; WT) and (E; *mpv17^a9/a9^*); and exposed neuromasts with disrupted morphology are shown in (C; WT) and (F; *mpv17^a9/a9^*). **(G-H)** Quantification of intact synapses per hair cell. Each data point refers to the average number of intact synapses per hair cell in one neuromast. Synapse number is reduced in both wild type and *mpv17^a9/a9^* neuromasts (G; **P=0.0016 WT; **P=0.0014 *mpv17^a9/a9^*). This reduction was consistent in both normal and disrupted neuromasts (H; **P=0.0090 WT normal; **P=0.0079 WT disrupted). Scale bar: 5 μm.

Studies in mammals have shown that noise exposures modulate the sizes of inner hair cell pre- and postsynaptic components (Kim et al., 2019; Song et al., 2016). We have also previously demonstrated that mechanical overstimulation of the lateral line resulted in enlarged PSDs (Holmgren et al., 2021). We thus measured the relative volumes of presynaptic ribbons and PSDs in *mpv17^a9/a9^* larvae and wild type siblings following mechanical overstimulation. We observed no significant change in the sizes of presynaptic ribbons in either wild type or *mpv17^a9/a9^* exposed neuromasts immediately following exposure (Fig. 8 E; Kruskal-Wallis test P=0.6165). When we compared PSD volumes, we observed enlarged PSD in unexposed *mpv17^a9/a9^* control neuromasts relative to wild type (Fig. 8 C,F; Dunn’s multiple comparison test *P=0.0100). In mechanically overstimulated wild type neuromasts, there was a dramatic increase in PSD size relative to control (Fig. 8 B,F; Dunn’s multiple comparison test ****P<0.0001). By contrast, in exposed *mpv17^a9/a9^* neuromasts, there was a modest, nonsignificant increase in PSD size relative to control, and the increase was less dramatic than in wild type (Fig. 8 D,F; Dunn’s multiple comparison test P=0.1787). These results reveal *mpv17^a9/a9^* mutant hair cells possess somewhat enlarged PSDs under homeostatic conditions and undergo less dramatic changes in PSD size following mechanical overstimulation relative to wild type.

**Figure 8.**
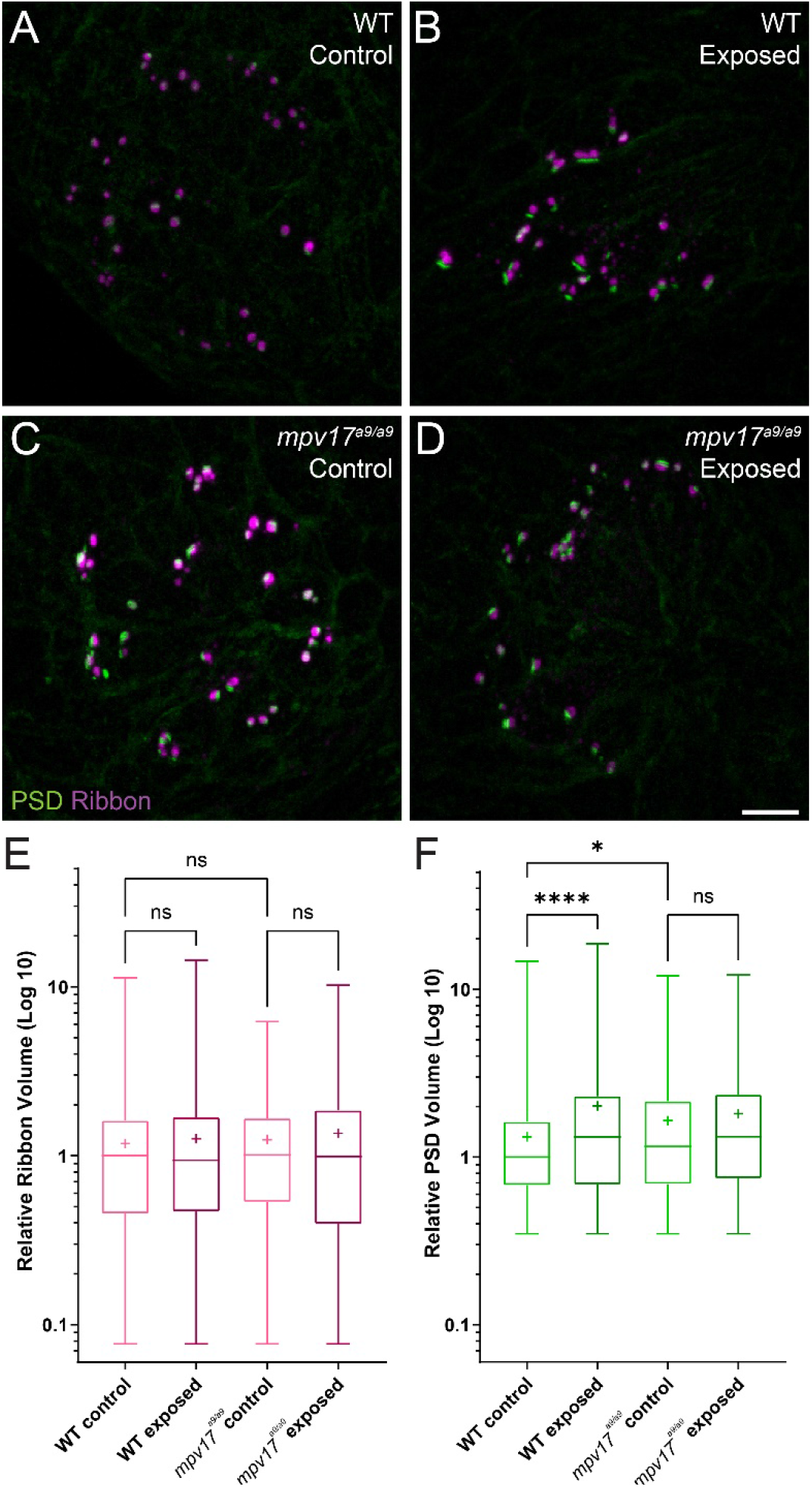
PSDs are enlarged in *mpv17^a9/a9^* neuromasts but do not significantly change upon mechanical overstimulation. **(A-D)** Maximum-intensity projections of confocal images of wild type (A,B) and *mpv17^a9/a9^* (C,D) neuromasts, either unexposed (A,C) or exposed to strong water current (B,D). **(E)** Quantification of presynaptic ribbon volume relative to unexposed wild type control. **(F)** Quantification of PSD volume relative to wild type control. PSDs are enlarged in unexposed *mpv17^a9/a9^* neuromasts and mechanically overstimulated wild type neuromasts, but there is no significant change in PSD size in mechanically overstimulated *mpv17^a9/a9^* neuromasts (****P<0.0001 WT exposed; *P=0.0100 *mpv17^a9/a9^* control). Scale bar: 5 μm.

## Discussion

### Effects of Mpv17 deficiency in zebrafish lateral-line hair cells

In this study, we have characterized the effects of Mpv17 deficiency on lateral-line hair cells, both under homeostatic conditions and in response to mechanical overstimulation. Mitochondrial homeostasis appeared to be disrupted, as we observed in *mpv17^a9/a9^* mutant hair cells elevated ROS and mitochondrial Ca^2+^, as well as hyperpolarized ΔΨ_m_. Mpv17-deficient hair cells showed reduced FM1-43 uptake and were more susceptible to neomycin-induced death. Following mechanical overstimulation, *mpv17^a9/a9^* neuromasts were more vulnerable to morphological disruption and hair-cell loss than wild type siblings, but showed similar degrees of de-innervation and synapse loss.

To our knowledge, while the transparent Casper mutant line of zebrafish is commonly used in research, this is the first investigation of the role of Mpv17 in lateral-line hair-cell mitochondrial dynamics. Mpv17 function has been studied more extensively in mammalian models (Binder, Weiher, Exner, & Kerjaschki, 1999; Meyer zum Gottesberge et al., 2001; Meyer zum Gottesberge et al., 2012; Muller et al., 1997). In mice, loss of Mpv17 results in severe defects in the kidney and inner ear. In humans, severe *MPV17* deficiency has been associated with a hepatocerebral form of mitochondrial DNA depletion syndrome which results in death due to liver failure at young ages, while less severe mutations have been linked to juvenile-onset peripheral neuropathy (Baumann et al., 2019; El-Hattab et al., 2018; Spinazzola et al., 2008). The phenotypes observed in *mpv17^a9/a9^* zebrafish are much milder than those reported in mammals, as homozygous mutants appear healthy and have normal lifespans. Paralogous genes in zebrafish may provide compensatory mechanisms for loss of Mpv17 (Glasauer & Neuhauss, 2014). Zebrafish have two *mpv17*-like genes, *mpv17l* and *mpv17l2,* and *mpv17l2* is upregulated in *mpv17^a9/a9^* fish (Krauss et al., 2013; Martorano et al., 2019). While it is known that *mpv17l2* is strongly expressed in the larval zebrafish liver (Thisse, 2004), it remains to be determined whether *mpv17l2* is expressed specifically in hair cells.

### Mitochondrial homeostasis in *mpv17^a9/a9^* hair cells

Mpv17-deficient hair cells showed elevated ROS and mitochondrial Ca^2+^, as well as a more negative ΔΨ_m_ (Fig. 1, 2). Mitochondrial Ca^2+^, ROS production, and ΔΨ_m_ are tightly linked in the cell (Adam-Vizi & Starkov, 2010; Brookes, Yoon, Robotham, Anders, & Sheu, 2004; Esterberg et al., 2014; Gorlach, Bertram, Hudecova, & Krizanova, 2015; Ivannikov & Macleod, 2013). ROS are generated in the mitochondria as a consequence of oxidative phosphorylation, which depends on a negative ΔΨ_m_. Negative ΔΨ_m_ is also maintained by removing protons from the mitochondrial matrix as electrons flow through the electron transport chain. A more negative ΔΨ_m_ thus results in more ROS (Kann & Kovacs, 2007; Zorov, Juhaszova, & Sollott, 2014). Mitochondrial Ca^2+^ regulates oxidative phosphorylation, so the elevated mitochondrial Ca^2+^observed in *mpv17^a9/a9^* hair cells we observed also contributes to increased ROS and ΔΨ_m_ (Brookes et al., 2004).

Similar hair-cell phenotypes were recently reported in *pappaa^p170^* mutant zebrafish: *pappaa^p170^* hair cells had elevated mitochondrial Ca^2+^, elevated ROS, and hyperpolarized mitochondria (Alassaf et al., 2019). Additionally, these mutants were more sensitive to neomycin-induced hair-cell loss, similar to what we observed in *mpv17^a9/a9^* neuromasts (Fig. 4). One notable difference between *pappaa^p170^ and mpv17^a9/a9^* mutant hair cells is that the significantly reduced FM1-43 uptake we observed in *mpv17^a9/a9^* mutants (Fig. 3) was not observed in *pappaa^p170^*, indicating that the driving force for cations through mechanotransduction channels is decreased in *mpv17^a9/a9^* but not *pappaa^p170^* mutants. Given that neomycin uptake is likely reduced in *mpv17^a9/a9^* mutant hair cells (Hailey et al., 2017), our observations support that mitochondrial disfunction is the main source of increased vulnerability to neomycin. In *pappaa^p170^* mutants, this susceptibility was rescued by treatment with the ROS scavenger mitoTEMPO, supporting that attenuating oxidative stress could rescue *mpv17^a9/a9^* susceptibility to neomycin. Additionally, a study defining neomycin-induced hair-cell death showed that mitochondrial Ca^2+^ uptake in zebrafish neuromast hair cells is a precursor to cell death, so it is possible that the elevated mitochondrial Ca^2+^ observed in *mpv17^a9/a9^* hair cells may contribute to the heightened sensitivity (Esterberg et al., 2014). This study also demonstrated that ΔΨ_m_ plays a role in neomycin ototoxicity such that partially depolarizing the mitochondria with sub-lethal levels of FCCP offered a protective effect. Cumulatively, our observations in *mpv17^a9/a9^* hair cells provide additional support to the idea that mitochondrial disfunction underlies enhanced susceptibility to neomycin-induced hair-cell death, and that drugs targeting ROS or that uncouple mitochondrial phosphorylation may provide therapeutic protection.

In larvae exposed to strong water current stimulus, *mpv17^a9/a9^* neuromasts displayed mechanical disruption of morphology more frequently than wild type siblings exposed to the same stimulus (Fig. 5). We also observed, in disrupted neuromasts, a significant decrease in hair-cell number which was exacerbated in *mpv17^a9/a9^* mutants (Fig. 6). In our previous study, we reported intact mechanotransduction was not required for stimulus-induced displacement of neuromasts, indicating this observed phenotype is a result of mechanical injury (Holmgren et al., 2021). While it is unclear why *mpv17^a9/a9^* neuromasts are more susceptible to mechanically induced displacement, it is possible that mitochondrial disfunction in hair cells of disrupted *mpv17^a9/a9^* neuromasts sensitizes them to mechanically induced death. It would be interesting to know whether pharmacological manipulation of ΔΨ_m_ could affect a neuromast’s vulnerability to mechanical injury and hair-cell loss. As *mpv17^a9/a9^* hair cells also showed elevated ROS (Fig. 1), oxidative stress could also play a role in morphological disruption and hair-cell loss. Further examination of ROS levels in mechanically overstimulated hair cells will be important to fully understand these mechanisms.

### Mechanical overstimulation and hair-cell synapse loss

It has been established that moderate noise exposure results in a loss of hair-cell synapses in the cochlea, but the mechanisms underlying noise-induced synapse loss are not completely understood (Kujawa & Liberman, 2009). Recent studies have defined roles of mitochondrial Ca^2+^ in synaptic maintenance and noise-induced synapse loss (Wang et al., 2018; Wong et al., 2019). One mechanism by which Ca^2+^ is taken up by mitochondria is via the mitochondrial Ca^2+^ uniporter (MCU) (Wong et al., 2019). Both inhibition of MCU by Ru360 and genetic deletion of MCU have been shown to prevent hair-cell synapse loss in noise-exposed mice (Wang et al., 2018). These observations support that mitochondrial Ca^2+^ uptake plays a critical role in noise-induced synapse loss. We have shown here that *mpv17^a9/a9^* hair cells have elevated mitochondrial Ca^2+^ levels, as measured by the genetically encoded mitochondrial Ca^2+^indicator MitoGCaMP3 (Fig. 1) (Esterberg et al., 2014). It would therefore be unsurprising if elevated mitochondrial Ca^2+^ in *mpv17^a9/a9^* hair cells contributed to more severe hair-cell synapse loss following mechanical overstimulation. However, here we observed similar degrees of synapse loss in both wild type and *mpv17^a9/a9^* stimulus exposed neuromasts (Fig. 8), suggesting an alternate mechanism is at play.

It has recently been shown that *Vglut3^−/−^* null mutant mice do not lose hair-cell synapses following noise exposure, supporting a role for synaptic transmission in noise-induced synapse loss (Kim et al., 2019). Additionally, in our previous study of mechanical injury in the zebrafish lateral line, we observed significantly more severe mechanically-induced hair-cell synapse loss in fish when glutamate clearance from the synapse was pharmacologically blocked, suggesting synapse loss can be exacerbated by excess glutamate in the synaptic cleft (Holmgren et al., 2021). Our results here show that *mpv17^a9/a9^* hair cells have reduced FM1-43 uptake indicating reduced hair-cell transduction (Fig. 3), which suggests synaptic transmission may also be impaired in *mpv17^a9/a9^* hair cells. It is possible that there is a balancing act in *mpv17^a9/a9^* hair cells, with elevated mitochondrial Ca^2+^ potentially tipping the scales toward more severe synapse loss, yet counter balanced by reduced hair-cell activity providing protection from glutamate excitotoxicity. Further studies will be necessary to determine the relative contribution of mitochondrial activity to mechanically induced hair-cell synapse loss.

### Conclusion

We have shown here that mitochondrial homeostasis is disrupted in *mpv17^a9/a9^* hair cells of the zebrafish lateral line. Consequently, *mpv17^a9/a9^* neuromast hair cells are more vulnerable to neomycin- and mechanically-induced hair-cell death, but are not more susceptible to synapse loss from overstimulation. This model will be useful for future studies examining not only the relative contributions of mitochondrial function to hair-cell damage, but also the roles of mitochondrial homeostasis in subsequent repair following damage.

## Declaration of interests

The authors declare no competing financial or non-financial interests.

## Acknowledgments

This work was supported by the National Institute on Deafness and Other Communication Disorders R01DC016066 (L.S.). We would like to thank Valentin Militchin for engineering support.

## Materials and Methods

### Ethics statement

Experimental procedures were performed with approval from the Washington University School of Medicine Institutional Animal Care and Use Committee and in accordance with NIH guidelines for use of zebrafish.

### Zebrafish

All zebrafish experiments and procedures were performed in accordance with the Washington University Institutional Animal Use and Care Committee. Adult zebrafish were raised under standard conditions at 27-29°C in the Washington University Zebrafish Facility. Embryos were raised in incubators at 28°C in E3 media (5 mM NaCl, 0.17 mM KCl, 0.33 mM CaCl2, 0.33 mM MgCl2; (Nüsslein-Volhard & Dahm, 2002) with a 14 h:10 h light:dark cycle. After 4 dpf, larvae were raised in 100-200 ml E3 media in 250-ml plastic beakers and fed rotifers daily. Sex of the animal was not considered for this study because sex cannot be determined in larval zebrafish.

The transgenic lines *TgBAC(neurod1:EGFP)* and *Tg(myo6b:mitoGCaMP3)* were used in this study (Esterberg et al., 2014; Obholzer et al., 2008). Fluorescent transgenic larvae were identified at 3-5 days post-fertilization (dpf) under anesthesia with 0.01% tricaine in E3 media. The *TgBAC(neurod1:EGFP)* and *Tg(myo6b:mitoGCaMP3)* lines were crossed into Casper compound mutants (*mitfa^w2/w2^, mpv17^a9/a9^*) (White et al., 2008). Homozygous *mpv17^a9/a9^* mutants were identified at 3-5 dpf by phenotype under a brightfield dissecting microscope based on the severe reduction of iridophores in the eyes (D’Agati et al., 2017).

### Experimental apparatus

This experimental device was previously described in (Holmgren et al., 2021). In brief, 6-well plates containing larvae were clamped to a custom magnesium head expander (Vibration & Shock Technologies, Woburn, MA) on a vertically-oriented Brüel+Kjær LDS Vibrator, V408 (Brüel and Kjær, Naerum, Denmark). The apparatus was housed in a custom sound-attenuation chamber. An Optiplex 3020 Mini Tower (Dell) with a NI PCIe-6321, X Series Multifunction DAQ (National Instruments) running a custom stimulus generation program (modified version of Cochlear Function Test Suite) was used to relay the stimulus signal to a Brüel+Kjær LDS PA100E Amplifier that drove a controlled 60 Hz vibratory stimulus along the plate’s dorsoventral axis (vertically). Two accelerometers (BU-21771, Knowles, Itasca, IL) were mounted to the head expander to monitor the vertical displacement of the plate. The output of the accelerometers was relayed through a custom accelerometer amplifier (EPL Engineering Core). A block diagram for the EPL Lateral Line Stimulator can be found here: https://www.masseyeandear.org/research/otolaryngology/eaton-peabody-laboratories/engineering-core.

### Mechanical overstimulation of zebrafish larvae

Larval zebrafish were exposed to strong water current as previously described (Holmgren et al., 2021). At 7 dpf, free-swimming zebrafish larvae were placed in untreated 6-well plates (Corning, Cat# 3736; well diameter: 34.8 mm; total well volume: 16.8 ml) with 6 ml E3 per well, pre-warmed to 28°C. Up to 20 larvae were placed in each well. Individual wells were sealed with Glad^®^ Press ‘n Seal plastic food wrap prior to placing the lid on the plate. An additional metal plate was fitted to the bottom of the multi-well dish to fill a small gap between the bottoms of the wells and the head expander.

Exposures (stimulus parameters: 60 Hz, 46.2 ± 0.3 m/s2) were conducted at room temperature (22-24°C) up to 2 hours after dawn. Exposure consisted of 20 minutes of stimulation followed by a 10-minute break and 2 hours of uninterrupted stimulation. During the entire duration of exposure, unexposed control fish were kept in the same conditions as noise-exposed fish i.e. placed in a multi-well dish and maintained in the same room as the exposure chamber. After exposure, larvae were immediately fixed for histology.

### Pharmacology

To assess hair-cell sensitivity to aminoglycosides, 5-6 dpf larvae were exposed to 10 μM neomycin (Sigma) in E3 for 30 minutes at 28°C. Larvae were then rinsed in E3 and allowed to recover for 2 hours at 28°C before being fixed for immunohistochemical labeling of hair cells.

To verify that entry of MitoTracker dyes into hair cells was not dependent on mechanotransduction, 7 dpf larvae were exposed to 5 mM BAPTA (Invitrogen) in E3 for 10 minutes, then rinsed in E3. MitoTracker probes were calibrated by treating fish with cyclosporin A (TCI America). Larvae were exposed to 200 nM cyclosporin A alone in E3 with 0.1% DMSO for 5 minutes prior to co-exposure with MitoTracker probes and drug for 30 minutes.

### Live hair-cell labeling

Hair cell nuclei were specifically labeled by briefly exposing free-swimming larvae to 4’,6-Diamidino-2-Phenylindole (DAPI; Invitrogen/ThermoFisher) diluted to 2.5 μg/ml in E3. Prior to exposure to mechanical stimulation or incubation with other indicators, larvae were exposed to DAPI working solution for 4 minutes, then rinsed 3 times in E3.

CellROX Green (Invitrogen) was used to quantify levels of ROS in hair cells. Larvae were exposed to 5 μM CellROX Green in E3 for 30 minutes at 28°C, protected from light. Larvae were then rinsed twice in E3 and prepared for live imaging. Larvae were maintained in the dark prior to imaging.

The fixable vital dye FM1-43FX (Invitrogen) was used to quantify hair-cell mechanotransduction as previously described (Holmgren et al., 2021). In brief, 7 dpf larvae were exposed to 3 μM FM1-43FX in E3 for 20 seconds then rinsed in E3. Larvae were then fixed (4% paraformaldehyde, 4% sucrose, 150 μM CaCl_2_ in 0.1 M phosphate buffer) overnight at 4°C and mounted on slides with elvanol (13% w/v polyvinyl alcohol, 33% w/v glycerol, 1% w/v DABCO (1,4 diazobicylo[2,2,2] octane) in 0.2 M Tris, pH 8.5).

We chose to use MitoTracker Red CMXRos and MitoTracker Deep Red (Invitrogen) to measure mitochondrial membrane potential because they are well retained after fixation (Mot et al., 2016; Pendergrass et al., 2004). 7 dpf larvae were exposed to 500 nM MitoTracker Red CMXRos and 500 nM MitoTracker Deep Red concurrently in E3 for 30 minutes at 28°C, protected from light. Larvae were then rinsed in E3, fixed in 4% paraformaldehyde in PBS overnight at 4°C, and mounted on slides with elvanol.

### Whole-mount immunohistochemistry

Larvae were briefly sedated on ice and fixed (4% paraformaldehyde, 4% sucrose, 150 μM CaCl2 in 0.1 M phosphate buffer) for 5 hours at 4°C. Larvae were then permeabilized in ice-cold acetone, blocked (2% goat serum, 1% bovine serum albumin, 1% DMSO in PBS), and incubated with primary antibodies diluted in blocking buffer. The following commercial primary antibodies were used in this study: GFP (1:500; Aves Labs, Inc; Cat# GFP-1020), Parvalbumin (1:2000; Thermo Fisher; Cat# PA1-933), Parvalbumin (1:2000; Abcam; Cat# ab11427), Parvalbumin (1:500; Sigma-Aldrich; Cat# P3088), MAGUK (K28/86; 1:500; NeuroMab, UC Davis; Cat# 75-029), CtBP (1:2000; Santa Cruz Biotechnology Cat# sc-55502. Custom affinity-purified antibody generated against Danio rerio Ribeye b (mouse IgG2a; 1:2000) (Sheets, Trapani, Mo, Obholzer, & Nicolson, 2011). Following primary antibody incubation, larvae were washed and incubated with diluted secondary antibodies conjugated to Alexa Fluor 488, Alexa Fluor 546, Dylight 549, Alexa Fluor 555, and Alexa Fluor 647 (Invitrogen). Larvae were then counterstained with DAPI (Sigma) and mounted on slides with elvanol.

### Live imaging

Live imaging of zebrafish larvae was performed as previously described (Holmgren et al., 2021). In brief, zebrafish larvae were anesthetized with 0.01% tricaine in E3, then mounted lateral-side up on a thin layer of 1.5-2% low-melt agarose in a tissue culture dish with a cover-glass bottom (World Precision Instruments) and covered in E3 media. Z-stack images with a z step of 1 μm (CellROX) or 0.5 μm (mito-GCaMP3) were acquired via an ORCA-Flash 4.0 V3 camera (Hamamatsu) using a Leica DM6 Fixed Stage microscope with an X-Light V2TP spinning disc confocal (60 micron pinholes) and a 63x/0.9 N.A. water immersion objective. Z-acquisition parameters w/ X-light spinning disc: 488 laser “20% power”, 150 ms per frame. The perimeter of each neuromast for subsequent analysis was captured using differential interference contrast imaging following fluorescent image acquisition. Image acquisition was controlled by MetaMorph software.

### Confocal imaging of fixed specimens

Fixed sample images were acquired using an LSM 700 laser scanning confocal microscope with a 63x 1.4 NA Plan-Apochromat oil-immersion objective (Carl Zeiss). Confocal stacks were collected with a z step of 0.3 μm over 7-10 μm with pixel size of 100 nm (x-y image size 51 x 51 μm). Acquisition parameters were established using the brightest control specimen such that just a few pixels reached saturation in order to achieve the greatest dynamic range in our experiments. For quantitative measurements such as particle volume or fluorescence intensity, parameters including gain, laser power, scan speed, dwell time, resolution, and zoom, were kept consistent between comparisons.

### Confocal image processing and analysis

Analysis was performed on blinded images. Digital images were processed using ImageJ software (Schneider, Rasband, & Eliceiri, 2012). To correct for drift in the z direction, images were adjusted using StackReg when appropriate (Thevenaz, Ruttimann, & Unser, 1998). Subsequent image processing for display within figures was performed using Illustrator software (Adobe).

To measure volume of synaptic puncta, raw images containing single immunolabel were subtracted for background using a 20-pixel rolling ball radius. Whole neuromasts were delineated based on Parvalbumin label in maximum-intensity projections using the freehand selection and “synchronize windows” tools. Puncta were defined as regions of immunolabel with pixel intensity above a determined threshold: threshold for Ribeye label was calculated using the Isodata algorithm (Ridler, 1978) on maximum-intensity projections, threshold for MAGUK label was calculated as the product of 7 times the average pixel intensity of the whole NM region in a maximum-intensity projection. Particle volume was measured using the 3D object counter using a lower threshold and minimum particle size of 10 voxels (Bolte & Cordelieres, 2006). To normalizes for differences in intensity across experimental trials, volume measurements were normalized to the median wild type control volume for each trial for each individual channel. Intact synapses were manually counted and defined as colocalized or adjoining MAGUK and Ribeye or CtBP puncta. The number of intact synapses per hair cell was approximated by dividing the number of synapses by the number of hair cells in the neuromast.

To measure fluorescence intensity of indicators across whole neuromasts, images containing single channels were background-subtracted using a rolling ball radius of the following sizes: images containing celllROX label or mitoGCaMP3 (50-pixel), images containing FM1-43FX label (100-pixel), and images containing individual MitoTracker labels (200-pixel). Whole neuromasts were delineated based on DIC images (celllROX label and mitoGCaMP3) or hair cell specific DAPI label in maximum-intensity projections, and mean intensity of the indicator was measured. Measurements from each experimental trial were normalized to the wild type control median value.

### Statistical analysis

Statistical analyses were performed using Prism 9 (Graphpad Software Inc). Datasets were confirmed for normality using the D’Agostino-Pearson test when appropriate. Statistical significance between two groups was determined by an unpaired Student’s t test or a Mann–Whitney U test, depending on the normality of the dataset. Statistical significance between multiple groups with normal distributions was determined by one-way ANOVA and appropriate post-hoc tests, and statistical significance between multiple groups with non-normal distributions was determined by a Kruskal-Wallis test and appropriate post-hoc tests. For datasets dependent on multiple independent variables, statistical significance was determined using two-way ANOVA and appropriate post-hoc tests.

---

**Supplemental Figure 1.**
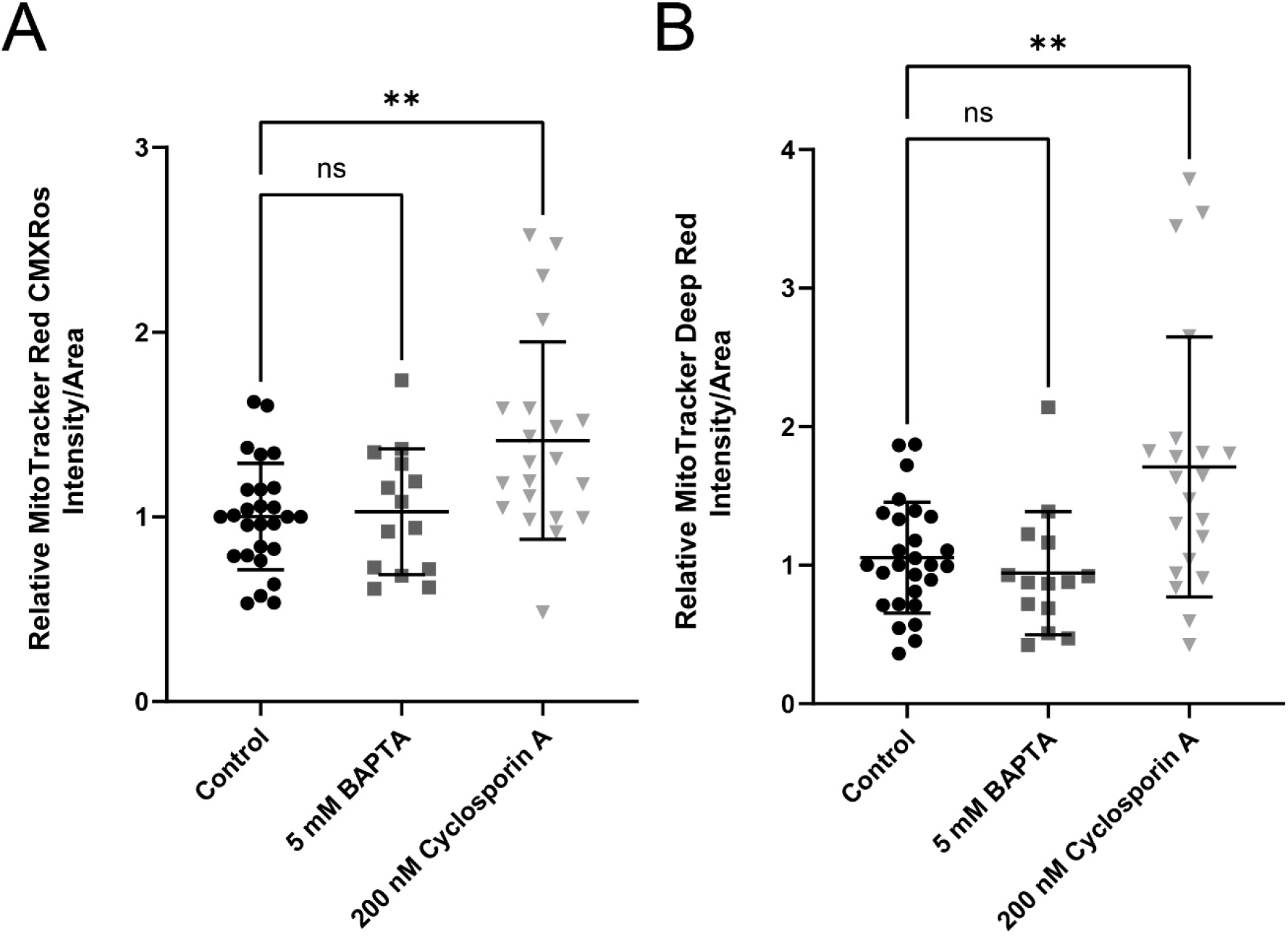
Validation of MitoTracker probes. Quantification of MitoTracker Red CMXRos (A) and MitoTracker Deep Red (B) fluorescence in WT neuromasts exposed to 200 nM cyclosporin A (to increase mitochondrial Δψm) or 5 mM BAPTA (to disrupt tip-links). Cyclosporin A treatment resulted in increased dye accumulation (**P=0.0022 MitoTracker Red CMXRos; **P=0.0025 MitoTracker Deep Red). BAPTA treatment did not significantly affect fluorescence, indicating hair-cell mechanotrasduction in not required for uptake of the dyes into hair cells.

## Notes

### Competing Interest Statement

The authors have declared no competing interest.

